# Age and sex-dependent patterns of gut microbial diversity in human adults

**DOI:** 10.1101/544270

**Authors:** Jacobo de la Cuesta-Zuluaga, Scott T. Kelley, Yingfeng Chen, Juan S. Escobar, Noel T. Mueller, Ruth E. Ley, Daniel McDonald, Shi Huang, Austin D. Swafford, Rob Knight, Varykina G. Thackray

**Affiliations:** Department of Microbiome Science, Max Planck Institute for Developmental Biology, 72076 Tübingen, Germany; Department of Biology, San Diego State University, San Diego, California, USA; Vidarium-Nutrition, Health and Wellness Research Center, Grupo Empresarial Nutresa, Medellin, Colombia; Department of Epidemiology, Johns Hopkins Bloomberg School of Public Health, Baltimore, MD, USA; Welch Center for Epidemiology, Prevention and Clinical Research, Johns Hopkins Medical Institutions, Baltimore, MD, USA; Department of Pediatrics, University of California San Diego, La Jolla, California, USA; Center for Microbiome Innovation, University of California, La Jolla, California, USA; Department of Computer Science, University of California San Diego, La Jolla, California, USA; Department of Bioengineering, University of California San Diego, La Jolla, California, USA; Department of Obstetrics, Gynecology and Reproductive Sciences, University of California San Diego, La Jolla, California, USA

**Author notes:** Contributed equally to this work (Jacobo de la Cuesta-Zuluaga, Scott T. Kelley).

**Keywords:** Diversity, Sex, Age, Microbiome, 16S rRNA amplicon

## Abstract

Gut microbial diversity changes throughout the human lifespan and is known to be affected by host sex. We investigated the association of age, sex and gut bacterial alpha diversity in three large cohorts of adults from four geographical regions: US and UK cohorts in the American Gut Project, and two independent cohorts of Colombians and Chinese. In three of the four cohorts, we observed a strong positive association between age and alpha diversity in young adults that plateaued after age 40. We also found pronounced sex-dependent differences in younger but not middle-aged adults, and women had higher alpha diversity than men. In contrast, no association of alpha diversity with age or sex was observed in the Chinese cohort. These associations were maintained after adjusting for cardiometabolic parameters in the Colombian cohort and antibiotic usage in the AGP cohort, suggesting that these factors do not affect the association of alpha diversity with age and sex. We also used a machine learning approach to predict individual age based on the gut microbiome. Consistent with our alpha diversity-based findings, women had significantly higher predicted age than men in the US and UK cohort, with a reduced difference above age 40. This was not observed in the Colombian cohort and only in the group of middle-age adults in the Chinese cohort. Together, our results provide new insights into the influence of age and sex on biodiversity of the human gut microbiota during adulthood while highlighting similarities and differences across diverse cohorts.

## Importance

Bacteria in the human gut play important roles in health and disease, and higher gut biodiversity has been linked to better health. Since gut bacteria may be pivotal in the development of microbial therapies, understanding the factors that shape gut biodiversity is of utmost interest. We performed large-scale analyses of the relationship of age and sex to gut bacterial diversity in adult cohorts from four geographic regions: US, UK, Colombia and China. In the US, UK and Colombia cohorts, bacterial biodiversity correlated positively with age in young adults, but plateaued around age 40 with no positive correlation in middle-aged adults. Young, but not middle-aged, adult women had higher gut bacterial diversity than men, a pattern confirmed via deep machine-learning. Interestingly, in the Chinese cohort, minimal associations were observed between gut biodiversity and age or sex. Our results highlight patterns of adult gut biodiversity and provide a framework for future research.

## Introduction

The human gut microbiota is a highly diverse ecosystem that is extremely variable amongst individuals (Lloyd-Price et al., 2017). This microbial community has been shown to play a key role in human health and disease (Gilbert et al., 2016). Since the gut microbiota may be pivotal to the development of microbial therapies, understanding factors that shape overall gut microbiota biodiversity over the different human life stages is of utmost interest.

There is increasing evidence suggesting that the host genes, gene expression patterns, environmental exposures, including medication and diet, and lifestyle factors play an important role in delimiting the boundaries of microbial diversity in the gut (Foster et al., 2017; McDonald et al., 2018). While a detailed longitudinal study of the interplay of each of these factors would be scientifically, logistically, and financially challenging, the chronological age of the host may be conceived as a proxy variable that represents the accumulation of these effects for a given individual. Several studies have reported a positive correlation between age and gut microbiota alpha diversity from birth to adulthood (Hopkins et al., 2002; Koenig et al., 2011; Mariat et al., 2009; Yatsunenko et al., 2012). Likewise, it has been shown that alpha diversity is maintained in old age until comorbidities contribute to its decline (Maffei et al., 2017). Another intriguing host-associated pattern identified in humans and rodents is the link between the gut microbiota and sex. Recent studies reported that women have higher microbial diversity than men and that sex influences the composition of the microbial community after puberty (Falony et al., 2016; Kozik et al., 2017; Markle et al., 2013; Sinha et al., 2018; Yatsunenko et al., 2012). These differences may contribute to the sexual dimorphism of autoimmune (Gomez et al., 2012; Markle et al., 2013; Yurkovetskiy et al., 2013) and neuro-immune diseases (Wallis et al., 2017, 2016). Therefore, it is key to consider the impact of inherent age and sex differences in different human populations to adequately discriminate changes and variations in the microbiome of individuals.

To better understand how age and sex of the host relate to the diversity of the gut microbiota during adulthood, we explored the association of these factors using data from individuals in three cross-sectional studies from four geographical origins including the citizen-science American Gut Project (AGP) comprised of individuals from the United States (US) and the United Kingdom (UK) (McDonald et al., 2018), a cohort of individuals from China (He et al., 2018) and a relatively smaller study with individuals from Colombia (de la Cuesta-Zuluaga et al., 2018).

## Materials and Methods

### Cohort description

Fecal samples were obtained from individuals in three independent cohorts from four geographical locations: i) The AGP dataset is composed of two cohorts with individuals from the UK (539 women and 397 men), and the US (1361 women and 1227 men) (Table 1); healthy participants self-reported age between 20 and 69, a body mass index (BMI) between 18.5 and 30 kg/m^2^, and no history of inflammatory bowel disease, diabetes, or antibiotic use in the past year. ii) A cohort of Chinese individuals (2772 women and 2191 men) aged 20 to 69, with BMI ranging from 18.5 to 30 kg/m^2^ and no antibiotic consumption reported one month prior to fecal sample collection; pregnant women, and hospitalized, disabled or critically-ill individuals were not included in the study. iii) A cohort of community-dwelling Colombians (226 women and 211 men), 20 to 62 years of age, enrolled in similar proportions according to: BMI, city of residence, and age range (20-40 and 41-62 years); underweight participants, pregnant women, individuals who had consumed antibiotics or antiparasitics in the three months prior to enrollment, and individuals diagnosed with neurodegenerative diseases, current or recent cancer (<1 year), and gastrointestinal diseases were excluded. Details on the data acquisition, quality assessment and processing of fecal samples from these three cohorts were previously described (de la Cuesta-Zuluaga et al., 2018; He et al., 2018; McDonald et al., 2018).

**Table 1.**
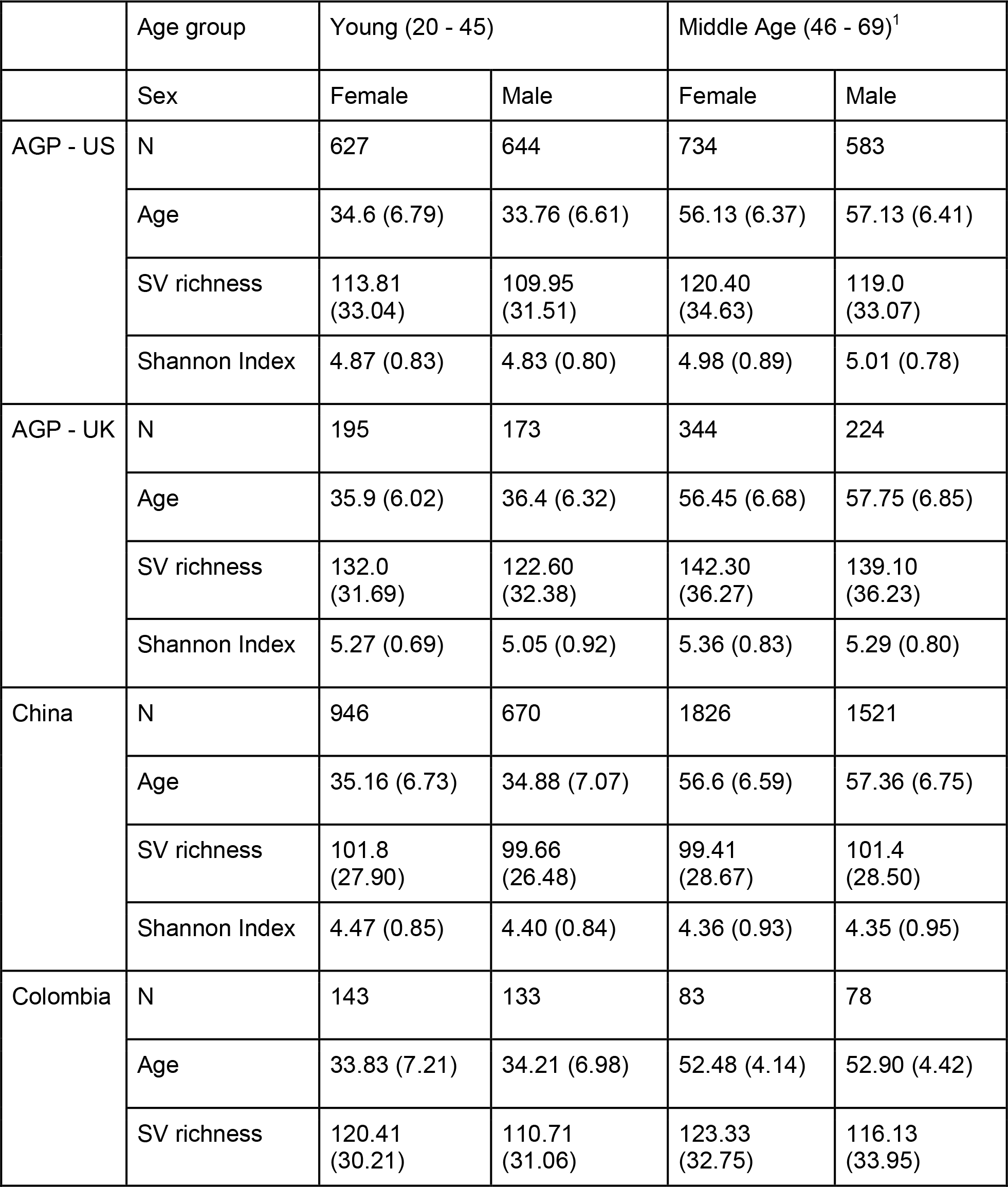
General characteristics of the participants of the included cohorts. Values given as mean (SD)

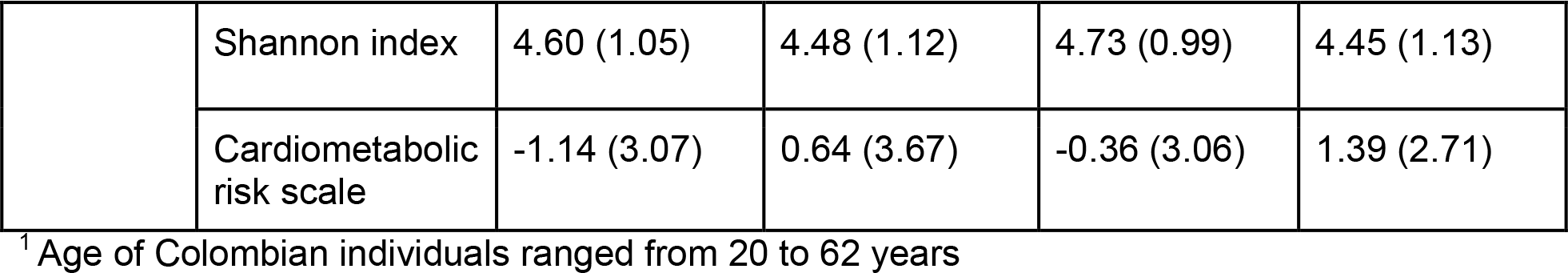

### 16S rRNA gene sequence processing

Amplicon sequences of all three cohorts were uniformly processed following the same procedures previously described (McDonald et al., 2018). Briefly, the V4 hypervariable region of the 16S rRNA gene was sequenced with the Illumina MiSeq platform. Raw sequences were clustered into sequence variants (SV) with deblur denoising (Amir et al., 2017) using QIIME 2 (Bolyen et al., 2018). Sequence counts were rarefied to 1250 reads per sample across all samples to mitigate uneven sequencing depth. Note, however, that sample collecting and DNA extraction methods differed between studies.

### Statistical analyses

SV richness and Shannon index were calculated using QIIME 2 and statistical analyses were performed using R v.3.4.3. The association of age and alpha diversity was measured with and without separate age groups by fitting linear models with linear splines with a knot at age 45 (lspline v.1.0 package of R) or simple linear models, respectively. We assessed the goodness of fit of these models by means of the Akaike’s information criterion (AIC). Next, scatter plots of each alpha diversity metric according to age were constructed and then separate LOESS curves for women and men were fitted using the ggplot2 v.3.0 package of R. Given the nonlinear association observed between alpha diversity and age, we subdivided the datasets into two separate age groups: 20 to 45 years (young adult) and 46 to 69 years (middle-aged adult), which were then used to fit linear models to test associations of age (as a continuous variable) and alpha diversity measures, stratified by sex.

Additionally, to account for possible confounder effects of antibiotic usage or cardiometabolic health of the subjects, we replicated the above analyses as follows. For the former, we replicated the analyses using a separate group of individuals of the AGP cohort from the US who had consumed antibiotics during the 6 months prior to their enrollment (283 women and 174 men). For the latter, we replicated the analyses adjusting the linear models for cardiometabolic risk in the Colombian cohort using a risk measure, which we termed cardiometabolic risk scale (Guzman-Castaneda et al., 2018). This was calculated using the sum of the z-scores of log-transformed waist circumference, triglycerides, insulin, diastolic blood pressure and high-sensitivity C reactive protein; positive values of the score are associated with increased cardiometabolic health risk.

Random Forest (RF) regression was used to regress relative abundances of SVs in the gut microbiota of healthy women and men adults against their chronological age in each dataset, using the R package randomForest and the following parameters: ntree = 18,000 and mtry = p/3, where p is the number of input features (SVs). The microbiota age model was first trained on the training dataset of female adults and was then applied to test the set of male adults, and vice versa. A smoothing spline function was fit between microbiota age and chronological age of the hosts for calculation of “relative microbiota age” of adults in the test sets to which the sparse model was applied. For a particular sample, the relative microbiota age was calculated as the difference between the “microbiota age of a focal adult” and the “microbiota age of interpolated spline fit of healthy women/men adults at the same chronological age”. We further employed Wilcoxon rank-sum test to compare the relative microbiota age between female and male groups in each of datasets using the R function Wilcox.test. To establish the sex difference in microbiota age, we subdivided the datasets into the above-defined age groups, and repeated the analyzes as described above in all age segments.

The code and data required to reproduce the statistical analyses is available at https://github.com/jacodela/microbio_aDiv.

## Results

Basic characteristics of individuals from the four cohorts are summarized in Table 1, stratified by sex and age group. To assess changes in alpha diversity with age during adulthood, we first performed linear regression with linear splines, establishing a knot at 45 years of age, and simple linear regressions in each cohort independently. We then evaluated the goodness of fit of each model using AIC, which indicated that changes in alpha diversity are better explained by distinguishing between young adults (20-45 years) and middle-aged adults (46-69 years). In the US, UK and Colombian cohorts, we observed a positive but non-linear association between alpha diversity measures and age, in both women and men. LOESS curves fit independently by sex showed an inflection point after age 40 in each of these cohorts (Fig. 1A-C). In contrast, we did not observe such a pattern in the Chinese cohort, in which alpha diversity displayed a slight decrease with age (Fig. 1D).

**Fig. 1.**
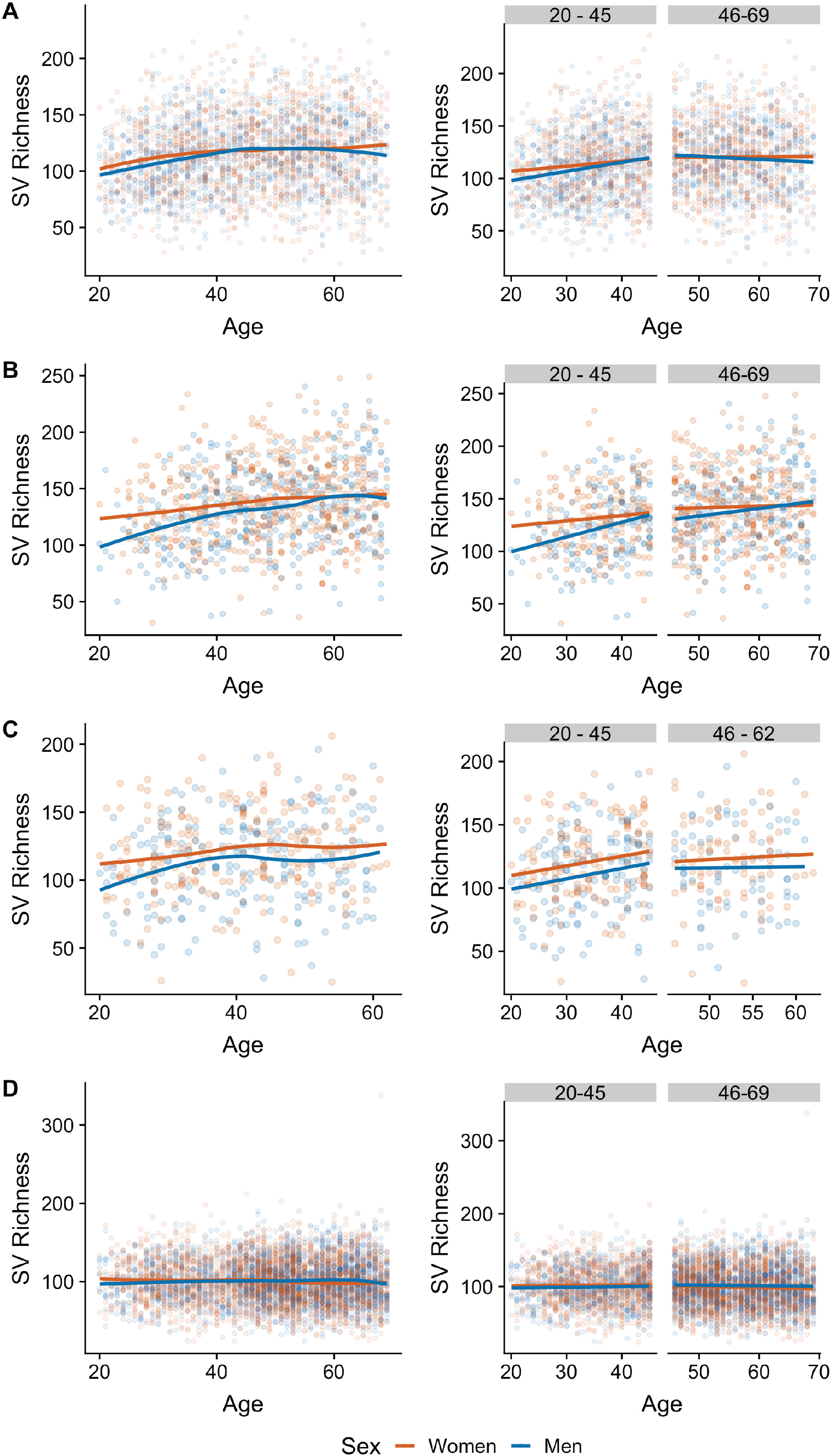

We then fit linear regression models to examine associations of microbial diversity, age and sex in the different age groups of each cohort independently. In both the US and UK cohorts, we observed a positive relationship between microbial richness and age for both sexes in young adults (US: p <0.001; UK: p < 0.001), but not in middle-aged adults (US: p = 0.404; UK: p = 0.111) (Fig. 1A-B). In addition, after adjusting for age we observed significant or borderline significant differences in overall SV richness between sexes in young adults (US: Δ_men - women_ = −3.3., p-value = 0.066; UK: Δ_men - women_ = −9.84, p-value = 0.003) but not middle-aged adults (US: Δ_men - women_ = −1.3, p-value = 0.48; UK: Δ_men - women_ = −3.7, p-value = 0.24). Similar results were observed when we assessed taxa evenness using the Shannon index (Fig. S1). Finally, we observed a significant interaction between age and sex with the Shannon index, but not SV richness, in young (Shannon US: p = 0.02; Shannon UK: p = 0.03; SV richness US: p= 0.15; SV richness UK: p= 0.09) but not middle-aged adults (p > 0.2 all comparisons).

To establish whether these trends were present in other populations, we then examined the association of alpha diversity with age and sex in the different age groups of cohorts from Colombia and China. Similar to the US and UK cohorts from the AGP, we identified a positive relationship between richness and age in the Colombian cohort in young adults of both sexes (p-value = 0.002) but not in middle-aged adults (p-value = 0.722) (Fig. 1C). Likewise, there was a significant difference in overall SV richness between the sexes in young adults (Δ_men - women_ = − 10.0; p-value = 0.006) but not in middle-aged adults (Δ_men - women_ = −7.3; p-value = 0.15). Potentially due to a smaller sample size, we did not find a significant interaction between age and sex on microbial diversity in the Colombian cohort for young (p = 0.91) or middle-aged adults (p = 0.81). In contrast to the US, UK and Colombian cohorts, we did not observe a relationship between biodiversity and age in young adults from the Chinese cohort (Fig. 1D). Men in the Chinese cohort tended to have lower SV richness compared to women as young adults, yet the difference was not significant (young adults: Δ_men - women_ = −4.46, p-value = 0.051; middle-aged adults: Δ_men - women_ = −4.08, p-value = 0.63).

Given that gut microbial diversity may be affected by factors such as antibiotic use or the cardiometabolic health of the host, we replicated the above analyses in cohorts in which we observed the patterns making use of publicly available metadata. To test whether the consumption of antibiotics affected the observed pattern, we performed the above analyses on a set of 457 individuals (283 women and 174 men) from the US cohort of the AGP that reported having consumed antibiotics in the 6 months prior to enrollment. As expected, we observed lower SV richness in these individuals compared to those that did not consume antibiotics. Additionally, our results indicated that the usage of antibiotics did not degrade the association of alpha diversity with age or sex in the AGP cohort (Fig. 2). Likewise, we replicated the analyses in the Colombian cohort after introducing a composite measure of the cardiometabolic health of the subjects as a covariate to the linear models; after we adjusted analyses for cardiometabolic health score, the observed patterns were similar (Fig. 3).

**Fig. 2.**
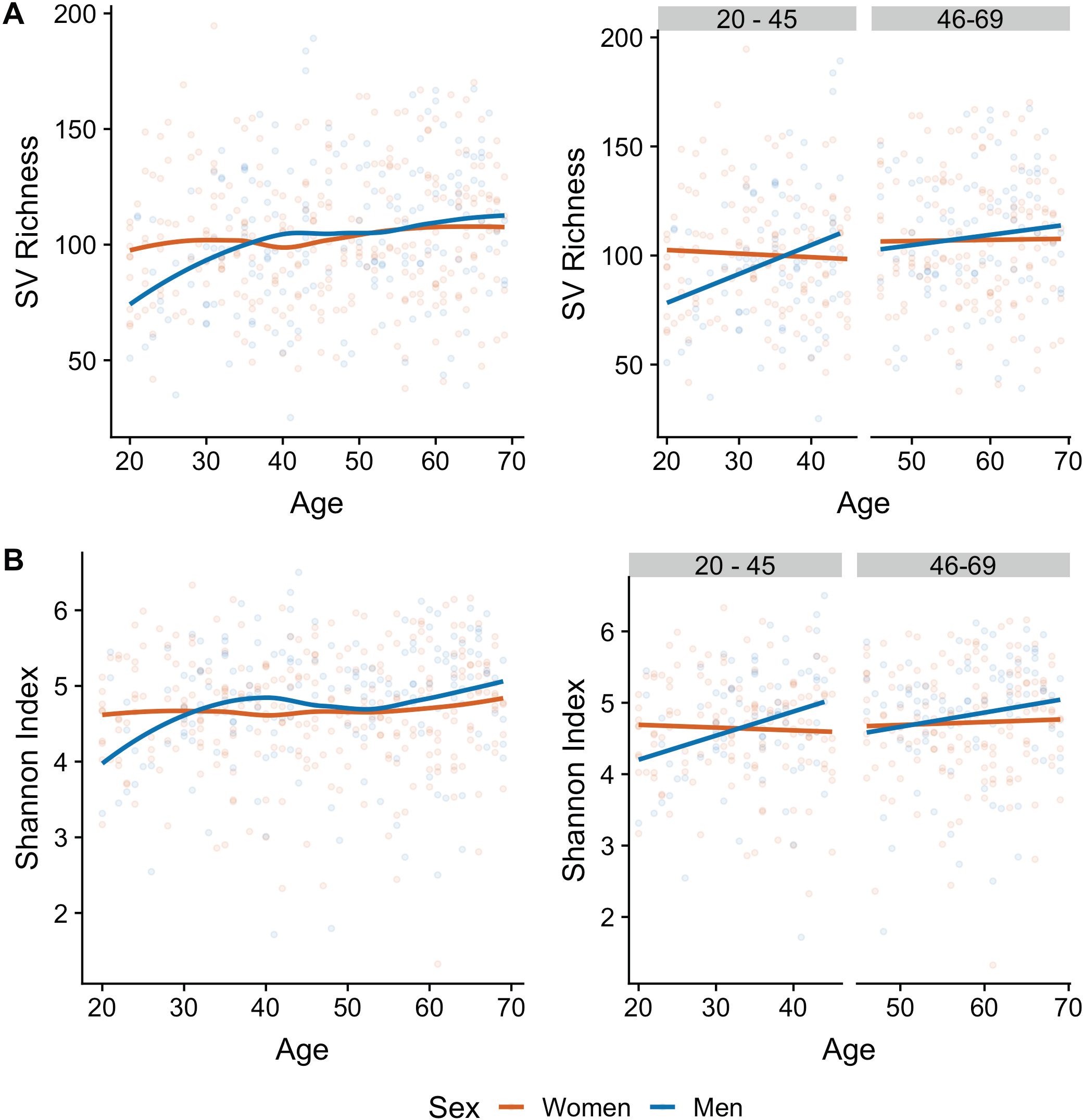

**Fig. 3.**
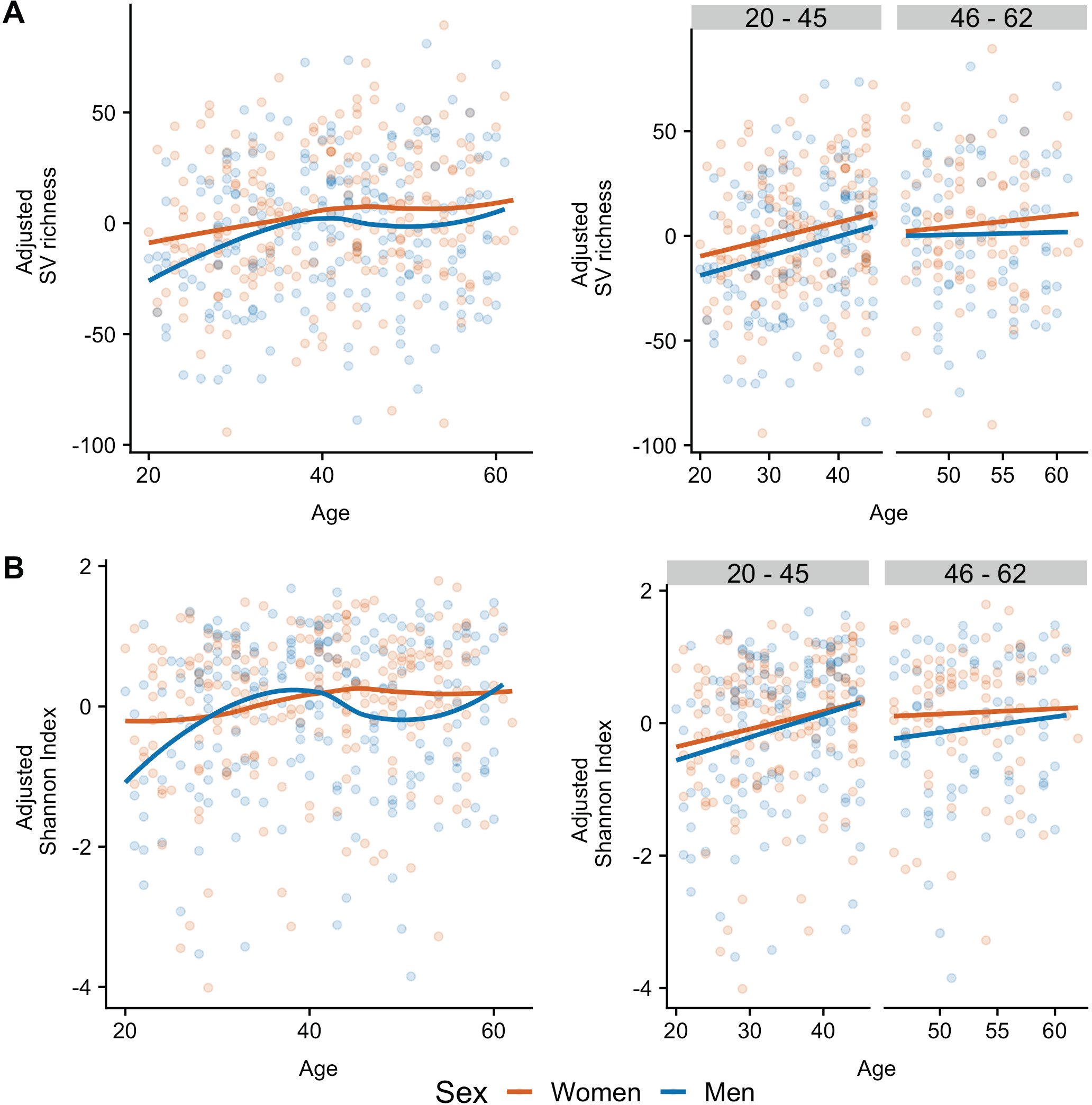

To examine whether similar age and sex-associated patterns in the gut microbiota would be observed applying an orthogonal method, we utilized a supervised machine learning approach using the composition of the gut microbiota of the subjects of the different populations. We subdivided each cohort by sex, determined the SVs shared by both groups, and used their relative abundances and the chronological age at time of sample collection of the host to fit a random forest (RF) regression model. Two models were built for women and men groups aged from 20-69 years; each trained using one sex and tested on the other. For each subject, we calculated the relative microbiota age as the difference between its microbiota age and the microbiota age of interpolated spline fit of an individual of the opposite sex at the same chronological age. Overall, our results indicate that the regressions can explain only 7-10 % of the variance in chronological age (from 20-69 years) of unrelated healthy adults from the US cohort.

We used 1494 shared SVs between women and men to build the RF model of the US cohort (Fig. 4A). We found that men exhibited lower relative microbiota age than women (p=6.237e-14, Wilcoxon rank-sum test; Fig. 4B, upper panels), suggesting sex may affect the adult gut microbial aging process. To validate this finding, we also trained a RF model in the men group and then applied to women (Fig. 4A, lower panels); we found women had higher microbiota age (p=2.467e-12, Wilcoxon rank-sum test. Fig. 4B, lower panels). To establish whether these trends were present in different age groups, we then examined the sex-dependent association of microbiota age in young and middle-aged separately. In the young group, we selected the 1311 shared SVs between both sexes to build the RF model for women and then applied it to predict the microbiota age of men. Compared to men, we found young women exhibited slightly higher relative microbiota age (p=0.0003225. Fig. 4C, D, upper panels). Similar results were observed when we assessed microbiota age in the middle-aged group, in which we used the 1601 shared SVs sexes to build the RF model as above. Microbiota age was higher in women compared to men (p=3.895e-15. Fig. 4C, D, upper panels). Furthermore, such sex difference in microbiota age were not affected when we applied men’s model to women data (Fig 4C, D, lower panels). Overall, our results indicate that the regressions can explain only 4.8-5.0 % of the variance in chronological age of unrelated healthy adults in the sex-stratified groups from the US cohort.

**Fig. 4.**
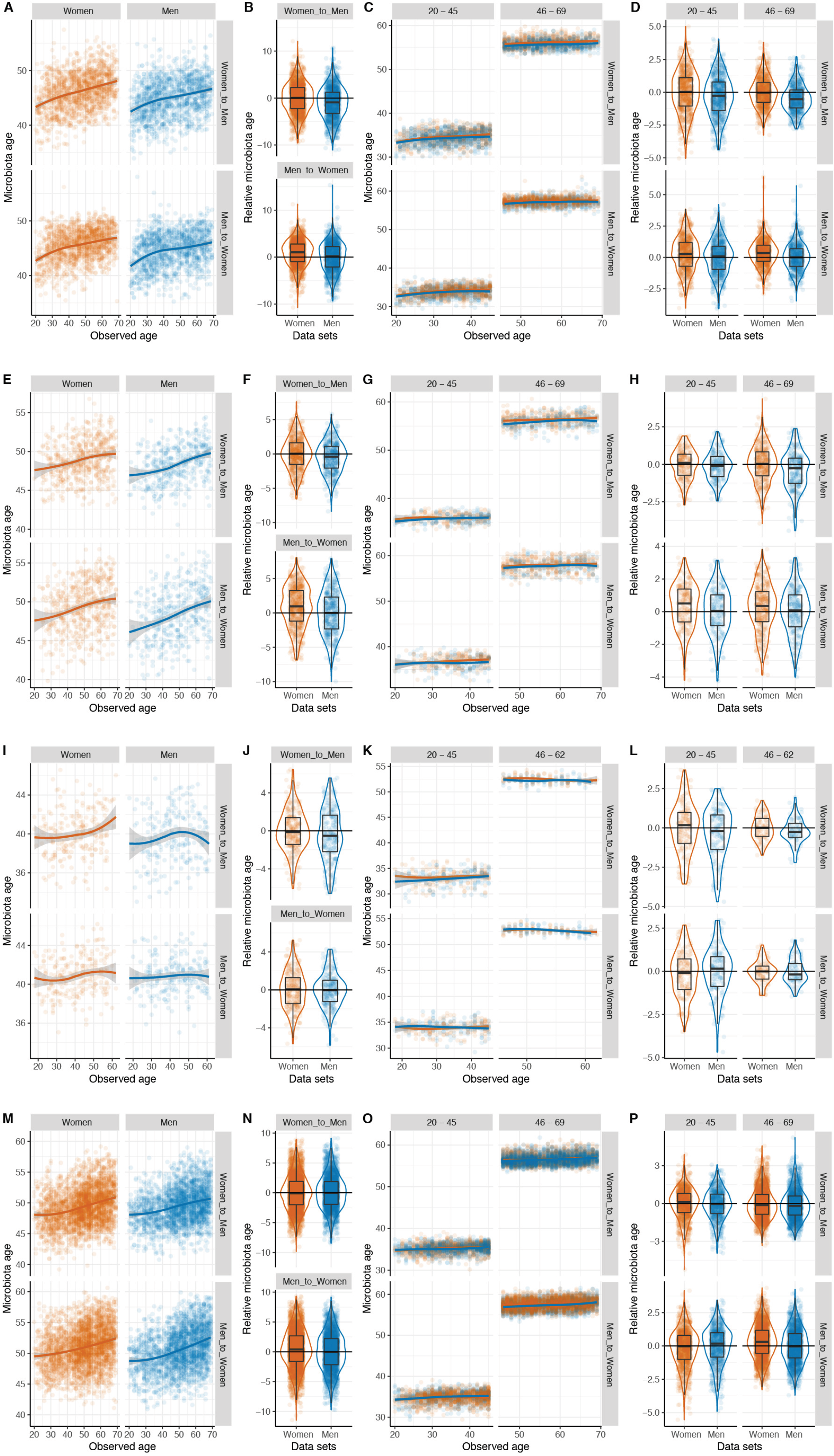

Likewise, in the UK cohort, we found higher microbiota age of women than that of men (from women to men: p= 0.001401; from men to women: p= 1.093e-05, Wilcoxon rank-sum test; Fig. 4E, F) using 1613 SVs in either women or men’s microbiota for building and applying RF models. In addition, we observed significant or borderline significant differences in relative microbiota age between sexes in young (from women to men: p= 0.1387; from men to women: p= 0.01376. Fig. 4G, H) and middle-aged (from women to men: p= 0.0001719; from men to women: p= 0.05378. Fig. 4G, H).

In the Colombian cohort we used 1074 SVs shared between sexes to build the RF model; similar yet non-significant trends were observed between microbiota age and sex in the non-stratified analyses (from women to men: p= 0.1103; from men to women: p= 0.9997, Wilcoxon rank-sum test; Fig. 4I, J), and in the young (from women to men: p= 0.1546; from men to women: p= 0.2519. Fig. 4K, L), and middle-aged groups (from women to men: p= 0.1435; from men to women: p= 0.8326. Fig. 4K, L). This is likely due to the smaller sample size of this cohort. We used 1279 SVs shared between sexes to build the RF models in the Chinese cohort, the association between microbiota age and sex in was not consistent when we cross-tested models (from women to men: p= 0.4248; from men to women: p= 2.422e-06. Fig. 4M, N). We did not observe significant associations in the young group (from women to men: p= 0.1249; from men to women: p= 0.01444. Fig. 4O, P), whereas in the middle-aged we observed sex-dependent difference of microbiota age and such differences are consistent in the cross-application of the models (from women to men: p= 0.03222; from men to women: p= 7.612e-12. Fig. 4O, P).

## Discussion

In this study, we analyzed the association of gut microbial alpha diversity with age and sex in three large cross-sectional cohorts encompassing four distinct human populations with a focus on gut bacteria of healthy adults. Our analyses indicate that age is positively associated with gut bacterial diversity in men and women, with greater diversity in women than men. Notably, this association occurs in young, but not middle-aged, adults. Consistent with these findings, predicted microbiota age varied based on sex with stronger effects seen in young adults. It is worth underscoring that while we did not observe these patterns in all studied cohorts, it was widespread, robust to technical differences, and the alpha diversity observations were not affected by the cardiometabolic health of the host nor disrupted by antibiotic consumption. These findings provide new insights into the development of the human gut microbiome during the course of life with regards to both age and sex, and emphasize the importance of including these factors as covariates in analyses of the human gut microbiota.

While the most dramatic change in gut microbiota diversity occurs in early childhood (Koenig et al., 2011; Yatsunenko et al., 2012), its increase in adulthood has also been reported (Odamaki et al., 2016). In the cohorts in which the pattern was present, we observed an increase in alpha diversity measures in young adults; however, this trend halted around age 40 (Figs 1, S1). This agrees with a previous report, in which no significant differences in alpha diversity were found between middle-aged and elderly subjects (Biagi et al., 2010). Interestingly, the relationship between age and diversity was also linked with sex. Multiple studies have reported differences in the diversity and composition of the gut microbiota between female and male mice, which appear to be associated with a sex bias in the incidence of specific diseases, such as type 1 diabetes (Markle et al., 2013; Yurkovetskiy et al., 2013), rheumatoid arthritis (Gomez et al., 2012), and anxiety (Bridgewater et al., 2017); sex-by-diet interactions have also been reported (Org et al., 2016). While differences in alpha diversity between males and females were reported in humans and mice, we showed that the association between sex and alpha diversity was stronger in young adults compared to middle-aged adults. In agreement with our results, no differences in alpha diversity were observed between women and men in a recent study in which the mean age of participants was 60 years (Haro et al., 2016).

One of the most intriguing findings was the difference in gut microbiota richness between the sexes in young adults. This sex-dependent discrepancy suggests that women may enter adulthood with a more diverse gut microbiota, which plateaus to the same levels in both sexes by approximately age 40. The establishment of different microbial communities in males and females may be mediated by sex hormones: female mice show a significant increase in alpha diversity during puberty (Kelley et al., 2016), and differences in the composition of the microbiota increase with age, but are eliminated by male castration (Yurkovetskiy et al., 2013). While little is known about the maturation of the human gut microbiota during puberty, we speculate that the differential hormonal milieu between women and men, and the earlier timing of puberty in women, may result in a more rapid diversification of the gut microbiota in women and that men only achieve the same level of diversification by middle age. Since our findings are based on cross-sectional data and cannot provide causal inference to test this hypothesis, future longitudinal studies that take into account factors such as steroid hormonal levels, age during different stages of puberty, contraceptive consumption, and pregnancy are needed to comprehend long-term trajectories of human gut microbial diversity.

While 3 of the 4 cohorts had an association between age, sex and microbial alpha diversity, the Chinese cohort did not (Figs 1, S1), indicating that these associations are a widespread but not universal feature of the human gut microbiota. The overall alpha diversity of this cohort, as measure by SV and the Shannon index, was lower than the other three cohorts in terms of SV. We also note that the exclusion criteria of this population was not the same as the other studies, with only a one-month antibiotic exclusion (versus 1 year) and no stated exclusion of participants with diabetes or inflammatory bowel disease (He et al. 2018).

The striking similarity among the US, UK and Colombian cohorts with regards to age and sex-dependent effects on microbial biodiversity arose despite different geographical origins, sample sizes and collection protocols of the studies. Moreover, we also found no apparent effect of antibiotic use (US or UK; Fig. 2) or cardiometabolic health (Colombia; Fig. 3) on the observed pattern in these cohorts, suggesting that the influence of age and sex on the microbiota may be similar in other ethnic and cultural groups, beyond the influence of cardiometabolic disease and antibiotics consumption. Nevertheless, similar large-scale population studies should be performed or reanalysed to determine the extent to which our results are generalizable to other populations, particularly in light of the Chinese cohort. Indeed, the contrast between the UK, US and Colombia cohorts compared to the Chinese cohort highlights the power of using large datasets and comparative analyses across cohorts to uncover subtle patterns and reveal novel insights not discernible in smaller studies. This is of critical importance given the plausibility of population-specific disease signatures of the microbiome (He et al., 2018).

## Acknowledgements

Data acquisition of the Colombian cohort was funded by Grupo Empresarial Nutresa, Dinámica IPS and EPS SURA. NTM was supported by the National Heart, Lung, and Blood Institute of the National Institutes of Health under Award Number K01HL141589, and by grants from the Mid-Atlantic Nutrition Obesity Research Center (P30DK072488) and the Foundation for Gender Specific Medicine. VGT was supported by the National Institute of Child Health and Human Development through a cooperative agreement as part of the National Centers for Translational Research in Reproduction and Infertility (P50 HD012303). STK and VGT received support from the Max Planck Institute for Developmental Biology in Tübingen. This work is supported by IBM Research AI through the AI Horizons Network and the UC San Diego Center for Microbiome Innovation. The content is solely the responsibility of the authors and does not necessarily represent the official views of the National Institutes of Health. Some authors of this work collaborate through the Microbiome & Health Network.

## References

Amir, A., McDonald, D., Navas-Molina, J.A., Kopylova, E., Morton, J.T., Zech Xu, Z., Kightley, E.P., Thompson, L.R., Hyde, E.R., Gonzalez, A., Knight, R., 2017. Deblur Rapidly Resolves Single-Nucleotide Community Sequence Patterns. mSystems 2.

Biagi, E., Nylund, L., Candela, M., Ostan, R., Bucci, L., Pini, E., Nikkïla, J., Monti, D., Satokari, R., Franceschi, C., Brigidi, P., De Vos, W., 2010. Through ageing, and beyond: gut microbiota and inflammatory status in seniors and centenarians. PLoS One 5, e10667.

Bolyen, E., Rideout, J.R., Dillon, M.R., Bokulich, N.A., Abnet, C., Al-Ghalith, G.A., Alexander, H., Alm, E.J., Arumugam, M., Asnicar, F., Bai, Y., Bisanz, J.E., Bittinger, K., Brejnrod, A., Brislawn, C.J., Titus Brown, C., Callahan, B.J., Caraballo-Rodríguez, A.M., Chase, J., Cope, E., Da Silva, R., Dorrestein, P.C., Douglas, G.M., Durall, D.M., Duvallet, C., Edwardson, C.F., Ernst, M., Estaki, M., Fouquier, J., Gauglitz, J.M., Gibson, D.L., Gonzalez, A., Gorlick, K., Guo, J., Hillmann, B., Holmes, S., Holste, H., Huttenhower, C., Huttley, G., Janssen, S., Jarmusch, A.K., Jiang, L., Kaehler, B., Kang, K.B., Keefe, C.R., Keim, P., Kelley, S.T., Knights, D., Koester, I., Kosciolek, T., Kreps, J., Langille, M.G.I., Lee, J., Ley, R., Liu, Y.-X., Loftfield, E., Lozupone, C., Maher, M., Marotz, C., Martin, B., McDonald, D., McIver, L.J., Melnik, A.V., Metcalf, J.L., Morgan, S.C., Morton, J., Naimey, A.T., Navas-Molina, J.A., Nothias, L.F., Orchanian, S.B., Pearson, T., Peoples, S.L., Petras, D., Preuss, M.L., Pruesse, E., Rasmussen, L.B., Rivers, A., Michael S Robeson, I.I., Rosenthal, P., Segata, N., Shaffer, M., Shiffer, A., Sinha, R., Song, S.J., Spear, J.R., Swafford, A.D., Thompson, L.R., Torres, P.J., Trinh, P., Tripathi, A., Turnbaugh, P.J., Ul-Hasan, S., van der Hooft, J.J.J., Vargas, F., Vázquez-Baeza, Y., Vogtmann, E., von Hippel, M., Walters, W., Wan, Y., Wang, M., Warren, J., Weber, K.C., Williamson, C.H.D., Willis, A.D., Xu, Z.Z., Zaneveld, J.R., Zhang, Y., Knight, R., Gregory Caporaso, J., 2018. QIIME 2: Reproducible, interactive, scalable, and extensible microbiome data science (No. e27295v1). PeerJ Preprints.

Bridgewater, L.C., Zhang, C., Wu, Y., Hu, W., Zhang, Q., Wang, J., Li, S., Zhao, L., 2017. Gender-based differences in host behavior and gut microbiota composition in response to high fat diet and stress in a mouse model. Sci. Rep. 7, 10776.

de la Cuesta-Zuluaga, J., Corrales-Agudelo, V., Velásquez-Mejía, E.P., Carmona, J.A., Abad, J.M., Escobar, J.S., 2018. Gut microbiota is associated with obesity and cardiometabolic disease in a population in the midst of Westernization. Sci. Rep. 8, 11356.

Falony, G., Joossens, M., Vieira-Silva, S., Wang, J., Darzi, Y., Faust, K., Kurilshikov, A., Bonder, M.J., Valles-Colomer, M., Vandeputte, D., Tito, R.Y., Chaffron, S., Rymenans, L., Verspecht, C., De Sutter, L., Lima-Mendez, G., Dhoe, K., Jonckheere, K., Homola, D., Garcia, R., Tigchelaar, E.F., Eeckhaudt, L., Fu, J., Henckaerts, L., Zhernakova, A., Wijmenga, C., Raes, J., 2016. Population-level analysis of gut microbiome variation. Science 352, 560–564.

Foster, K.R., Schluter, J., Coyte, K.Z., Rakoff-Nahoum, S., 2017. The evolution of the host microbiome as an ecosystem on a leash. Nature 548, 43–51.

Gilbert, J.A., Quinn, R.A., Debelius, J., Xu, Z.Z., Morton, J., Garg, N., Jansson, J.K., Dorrestein, P.C., Knight, R., 2016. Microbiome-wide association studies link dynamic microbial consortia to disease. Nature 535, 94–103.

Gomez, A., Luckey, D., Yeoman, C.J., Marietta, E.V., Berg Miller, M.E., Murray, J.A., White, B.A., Taneja, V., 2012. Loss of sex and age driven differences in the gut microbiome characterize arthritis-susceptible 0401 mice but not arthritis-resistant 0402 mice. PLoS One 7, e36095.

Guzman-Castaneda, S.J., Ortega-Vega, E.L., de la Cuesta-Zuluaga, J., Velasquez-Mejia, E.P., Rojas, W., Bedoya, G., Escobar, J.S., 2018. Gut microbiota composition explains more variance in the host cardiometabolic risk than genetic ancestry. bioRxiv.

Haro, C., Rangel-Zúñiga, O.A., Alcalá-Díaz, J.F., Gómez-Delgado, F., Pérez-Martínez, P., Delgado-Lista, J., Quintana-Navarro, G.M., Landa, B.B., Navas-Cortés, J.A., Tena-Sempere, M., Clemente, J.C., López-Miranda, J., Pérez-Jiménez, F., Camargo, A., 2016. Intestinal Microbiota Is Influenced by Gender and Body Mass Index. PLoS One 11, e0154090.

He, Y., Wu, W., Zheng, H.-M., Li, P., McDonald, D., Sheng, H.-F., Chen, M.-X., Chen, Z.-H., Ji, G.-Y., Zheng, Z.-D.-X., Mujagond, P., Chen, X.-J., Rong, Z.-H., Chen, P., Lyu, L.-Y., Wang, X., Wu, C.-B., Yu, N., Xu, Y.-J., Yin, J., Raes, J., Knight, R., Ma, W.-J., Zhou, H.-W., 2018. Regional variation limits applications of healthy gut microbiome reference ranges and disease models. Nat. Med.

Hopkins, M.J., Sharp, R., Macfarlane, G.T., 2002. Variation in human intestinal microbiota with age. Dig. Liver Dis. 34 Suppl 2, S12–8.

Kelley, S.T., Skarra, D.V., Rivera, A.J., Thackray, V.G., 2016. The Gut Microbiome Is Altered in a Letrozole-Induced Mouse Model of Polycystic Ovary Syndrome. PLoS One 11, e0146509.

Koenig, J.E., Spor, A., Scalfone, N., Fricker, A.D., Stombaugh, J., Knight, R., Angenent, L.T., Ley, R.E., 2011. Succession of microbial consortia in the developing infant gut microbiome. Proc. Natl. Acad. Sci. U. S. A. 108 Suppl 1, 4578–4585.

Kozik, A.J., Nakatsu, C.H., Chun, H., Jones-Hall, Y.L., 2017. Age, sex, and TNF associated differences in the gut microbiota of mice and their impact on acute TNBS colitis. Exp. Mol. Pathol. 103, 311–319.

Lloyd-Price, J., Mahurkar, A., Rahnavard, G., Crabtree, J., Orvis, J., Hall, A.B., Brady, A., Creasy, H.H., McCracken, C., Giglio, M.G., McDonald, D., Franzosa, E.A., Knight, R., White, O., Huttenhower, C., 2017. Strains, functions and dynamics in the expanded Human Microbiome Project. Nature 550, 61–66.

Maffei, V.J., Kim, S., Blanchard, E., 4th, Luo, M., Jazwinski, S.M., Taylor, C.M., Welsh, D.A., 2017. Biological Aging and the Human Gut Microbiota. J. Gerontol. A Biol. Sci. Med. Sci. 72, 1474–1482.

Mariat, D., Firmesse, O., Levenez, F., Guimarăes, V., Sokol, H., Doré, J., Corthier, G., Furet, J.-P., 2009. The Firmicutes/Bacteroidetes ratio of the human microbiota changes with age. BMC Microbiol. 9, 123.

Markle, J.G.M., Frank, D.N., Mortin-Toth, S., Robertson, C.E., Feazel, L.M., Rolle-Kampczyk, U., von Bergen, M., McCoy, K.D., Macpherson, A.J., Danska, J.S., 2013. Sex differences in the gut microbiome drive hormone-dependent regulation of autoimmunity. Science 339, 1084–1088.

McDonald, D., Hyde, E., Debelius, J.W., Morton, J.T., Gonzalez, A., Ackermann, G., Aksenov, A.A., Behsaz, B., Brennan, C., Chen, Y., DeRight Goldasich, L., Dorrestein, P.C., Dunn, R.R., Fahimipour, A.K., Gaffney, J., Gilbert, J.A., Gogul, G., Green, J.L., Hugenholtz, P., Humphrey, G., Huttenhower, C., Jackson, M.A., Janssen, S., Jeste, D.V., Jiang, L., Kelley, S.T., Knights, D., Kosciolek, T., Ladau, J., Leach, J., Marotz, C., Meleshko, D., Melnik, A.V., Metcalf, J.L., Mohimani, H., Montassier, E., Navas-Molina, J., Nguyen, T.T., Peddada, S., Pevzner, P., Pollard, K.S., Rahnavard, G., Robbins-Pianka, A., Sangwan, N., Shorenstein, J., Smarr, L., Song, S.J., Spector, T., Swafford, A.D., Thackray, V.G., Thompson, L.R., Tripathi, A., Vázquez-Baeza, Y., Vrbanac, A., Wischmeyer, P., Wolfe, E., Zhu, Q., American Gut Consortium, Knight, R., 2018. American Gut: an Open Platform for Citizen Science Microbiome Research. mSystems 3.

Odamaki, T., Kato, K., Sugahara, H., Hashikura, N., Takahashi, S., Xiao, J.-Z., Abe, F., Osawa, R., 2016. Age-related changes in gut microbiota composition from newborn to centenarian: a cross-sectional study. BMC Microbiol. 16, 90.

Org, E., Mehrabian, M., Parks, B.W., Shipkova, P., Liu, X., Drake, T.A., Lusis, A.J., 2016. Sex differences and hormonal effects on gut microbiota composition in mice. Gut Microbes 7, 313–322.

Sinha, T., Vich Vila, A., Garmaeva, S., Jankipersadsing, S.A., Imhann, F., Collij, V., Bonder, M.J., Jiang, X., Gurry, T., Alm, E.J., D’Amato, M., Weersma, R.K., Scherjon, S., Wijmenga, C., Fu, J., Kurilshikov, A., Zhernakova, A., 2018. Analysis of 1135 gut metagenomes identifies sex-specific resistome profiles. Gut Microbes 1–9.

Wallis, A., Butt, H., Ball, M., Lewis, D.P., Bruck, D., 2016. Support for the Microgenderome: Associations in a Human Clinical Population. Sci. Rep. 6, 19171.

Wallis, A., Butt, H., Ball, M., Lewis, D.P., Bruck, D., 2017. Support for the microgenderome invites enquiry into sex differences. Gut Microbes 8, 46–52.

Yatsunenko, T., Rey, F.E., Manary, M.J., Trehan, I., Dominguez-Bello, M.G., Contreras, M., Magris, M., Hidalgo, G., Baldassano, R.N., Anokhin, A.P., Heath, A.C., Warner, B., Reeder, J., Kuczynski, J., Caporaso, J.G., Lozupone, C.A., Lauber, C., Clemente, J.C., Knights, D., Knight, R., Gordon, J.I., 2012. Human gut microbiome viewed across age and geography. Nature 486, 222–227.

Yurkovetskiy, L., Burrows, M., Khan, A.A., Graham, L., Volchkov, P., Becker, L., Antonopoulos, D., Umesaki, Y., Chervonsky, A.V., 2013. Gender bias in autoimmunity is influenced by microbiota. Immunity 39, 400–412.

